# A chromosomal inversion may facilitate adaptation despite periodic gene flow in a freshwater fish

**DOI:** 10.1101/2021.12.02.470985

**Authors:** Matt J. Thorstensen, Peter T. Euclide, Jennifer D. Jeffrey, Yue Shi, Jason R. Treberg, Douglas A. Watkinson, Eva C. Enders, Wesley A. Larson, Yasuhiro Kobayashi, Ken M. Jeffries

## Abstract

Differences in genomic architecture between populations, such as chromosomal inversions, may play an important role in facilitating adaptation despite opportunities for gene flow. One system where chromosomal inversions may be important for eco-evolutionary dynamics are in freshwater fishes, which often live in heterogenous environments characterized by varying levels of connectivity and varying opportunities for gene flow. In the present study, reduced representation sequencing was used to study possible adaptation in *n* = 345 walleye (*Sander vitreus*) from three North American waterbodies: Cedar Bluff Reservoir (Kansas, USA), Lake Manitoba (Manitoba, Canada), and Lake Winnipeg (Manitoba, Canada). Haplotype and outlier-based tests revealed a putative chromosomal inversion that contained three expressed genes and was nearly fixed for alternate genotypes in each Canadian lake. These patterns exist despite several opportunities for gene flow between these proximate Canadian lakes, suggesting that the inversion may be important for facilitating adaptive divergence between the two lakes despite gene flow. Our study illuminates the importance of genomic architecture for facilitating local adaptation in freshwater fishes. Furthermore, our results provide additional evidence that inversions may facilitate local adaptation in many organisms that inhabit connected but heterogenous environments.

## Introduction

Differences in genomic architecture genomic architecture such as structural variants may help to facilitate local adaptation, and are therefore key mechanisms of eco-evolutionary processes (Dorant et al., 2020; Shi et al., 2021; Tigano & Friesen, 2016; Wellenreuther & Bernatchez, 2018). One type of structural variant, chromosomal inversions, are taxonomically widespread reversed regions of DNA that have been linked to both adaptation and speciation (reviewed in Wellenreuther & Bernatchez, 2018). Inversions can effectively protect large chromosomal regions containing hundreds of genes from recombination, facilitating local adaptation even in the face of high gene flow (Wellenreuther & Bernatchez, 2018). Most inversions found in previous studies have been relatively large because commonly used reduced representation sequencing methods can more easily detect large inversions (Wellenreuther & Bernatchez, 2018). However, inversions of any size may be important. According to a simulation study, intermediate-to-large inversions have been linked to local adaptation, but small inversions were preferentially fixed in directly beneficial mutation models and contributed the most to genome evolution in neutral and underdominant scenarios (Connallon & Olito, 2021). Among fishes, chromosomal inversions have largely been characterized in well studied groups of species such as representative salmonids (Arostegui et al., 2019; Leitwein et al., 2017; McKinney et al., 2020; Pearse et al., 2014) and gadids (Berg et al., 2017; Berg et al., 2016; Kirubakaran et al., 2016; Puncher et al., 2019; Sinclair-Waters et al., 2018; Sodeland et al., 2016), but little information exists on the importance of inversions in most taxa especially those inhabiting fresh water (Shi et al., 2021).

Freshwater fish often live in heterogenous environments characterized by varying levels of connectivity and therefore, varying opportunities for gene flow (Griffiths, 2015; Mushet et al., 2019). Gene flow tends to work in contrast to local adaptation (Lenormand, 2002) but nevertheless, evidence for local adaptation has been observed among different fish that use freshwater habitats (see Fraser et al., 2011 for a review of salmonids; Shi et al., 2021). While genomic architecture has been proposed to facilitate local adaptation (Tigano & Friesen, 2016), few studies test for this possibility in freshwater fishes (Shi et al., 2021). Freshwater habitats contribute a disproportionate richness in species diversity to global species diversity, given that only 0.01% of the planet’s total surface is fresh water (Balian et al., 2008). Fishes are a significant proportion of overall freshwater species diversity (approx. 10% of overall freshwater species and 70% of vertebrate freshwater species are fish; Balian et al., 2008). Therefore, an understanding of the mechanisms of local adaptation in freshwater fishes can contribute substantially to our overall understanding of local adaptation in heterogenous environments.

Walleye (*Sander vitreus*, but see Bruner, 2021) is a freshwater fish with a native range from the Northwest Territories (Canada) to Alabama (USA) (Hartman, 2009) and support important commercial and recreational fisheries (Fisheries and Oceans Canada, 2019, 2021). The economic importance of walleye underscores the need for research supporting the sustainable management of walleye fisheries, while its wide native distribution provides an opportunity to study the genomic basis of adaptation to a variety of freshwater environments. In addition, northern populations of walleye (such as those in Manitoba, Canada) were suspected to have been recently isolated into smaller lakes following the drainage of glacial Lake Agassiz approximately 7,000 years ago (Rempel & Smith, 1998; Stepien et al., 2009). This event may have resulted in walleye populations that remain closely related at neutral markers but show evidence of divergence at adaptive markers, such as in chromosomal inversions.

We sought to study how heterogenous environments with varying connectivity may have shaped contemporary walleye populations, and the potential for genomic architecture to facilitate adaptation despite opportunities for gene flow. Walleye from Lake Manitoba and Lake Winnipeg in Manitoba (Canada) were analyzed and compared with an outgroup from Cedar Bluff Reservoir in Kansas (USA) (Fig 1). Walleye in Lake Manitoba and Lake Winnipeg live in largely separated waterbodies with a shared river—Lake Manitoba drains eastward into Lake Winnipeg via the Fairford River, Lake St. Martin, and the Dauphin River (Fig 1). As a mobile species that undergoes broad seasonal migrations, historic gene flow may have been extensive between these Manitoba waterbodies (Backhouse-James & Docker, 2012; Munaweera Arachchilage et al., 2021; Thorstensen et al., 2020; Turner et al., 2021). In addition, historical flooding in 1882, 1902, 1904, 2011, and 2014 brought significantly increased waterflow eastward from Lake Manitoba toward Lake Winnipeg (Ahmari et al., 2016), and such flooding could have facilitated increased fish movement between the waterbodies. Last, walleye fry were opportunistically stocked in Lake Winnipeg from Lake Manitoba (Manitoba, Canada) between 1917-2002 (Manitoba Government, 2020). While intensive stocking complicated analyses of walleye ancestry in walleye of Minnesota and Wisconsin (USA) (Bootsma et al., 2020), stocking of walleye fry, which likely experience significant mortality, had little to no effect in South Dakota and Missouri (USA) (Fielder, 1992; Koppelman et al., 1992). Therefore, the genetic impacts of fry stocking in this system are likely minimal.

**Figure 1.**
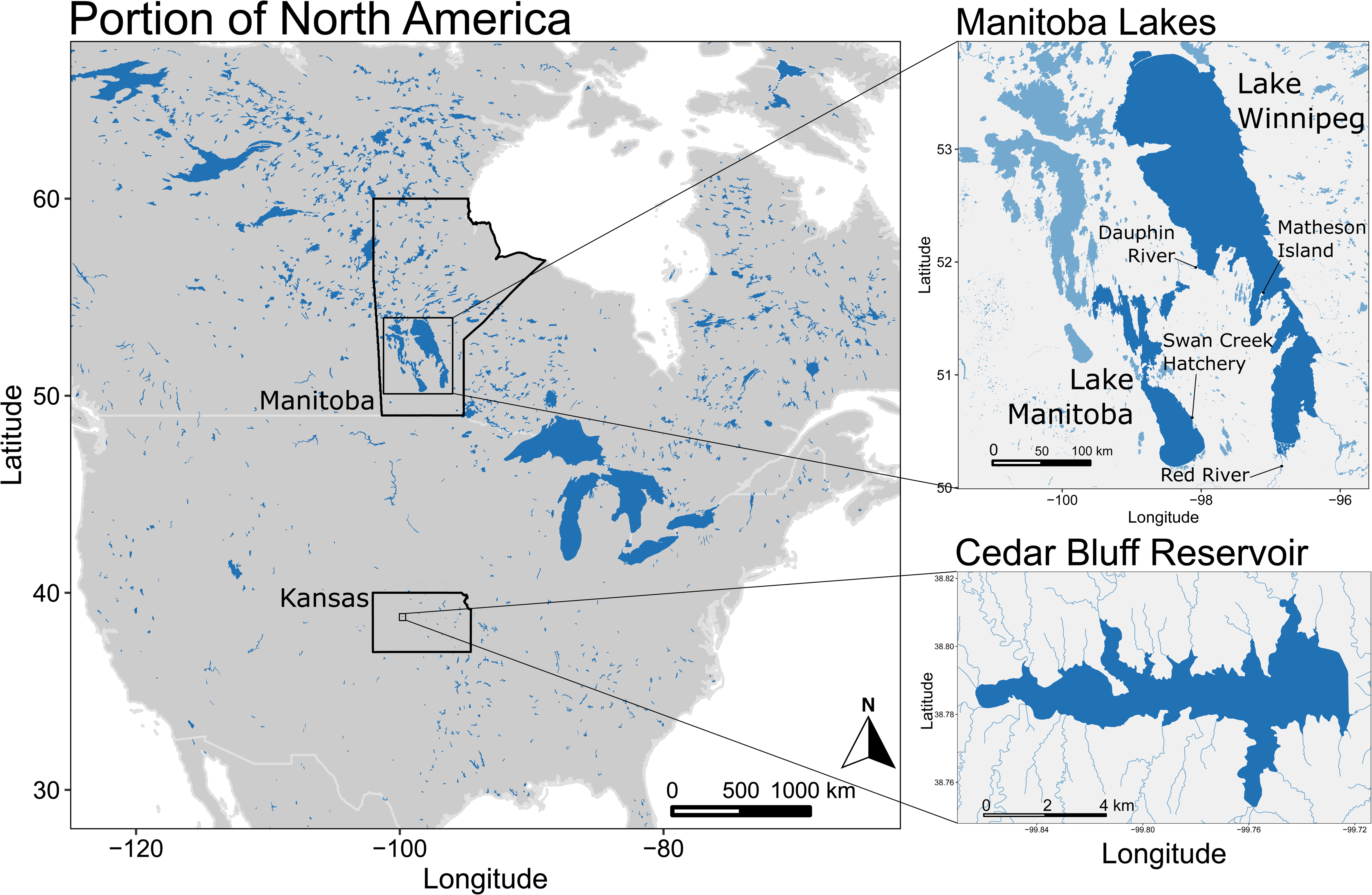
Map of waterbodies included in the present study. Cedar Bluff Reservoir, Kansas, USA, represents walleye (*Sander vitreus*) in an entirely stocked population of unknown origin. Lake Manitoba and Lake Winnipeg (Manitoba, Canada) walleye represent native populations, although Lake Winnipeg has been stocked by Lake Manitoba walleye throughout the 20^th^ century. In addition, floods in 1882, 1902, 1904, 2011, and 2014 increased waterflow from Lake Manitoba to Lake Winnipeg, and may have facilitated walleye gene flow.

Here, we employed Rapture (Ali et al., 2016), a reduced representation approach, to genotype *n* = 345 individuals from the three waterbodies described above. Population structure and demographic history were explored to provide context for signatures of selection. Simulations with EASYPOP were used to estimate historical migration rates between Lake Manitoba and Lake Winnipeg (Balloux, 2001). Candidate regions of adaptive variation were identified with XP-EHH, and outlier tests were used to confirm haplotype-based results. In addition, mRNA transcript expression in one candidate chromosomal inversion was assessed for Lake Winnipeg walleye to show the potential for the functional significance of genomic architecture in this system. The present study illuminates the importance of genomic architecture for facilitating local adaptation in freshwater fishes, and provides additional evidence that inversions may facilitate local adaptation in many organisms that inhabit connected but heterogenous environments.

## Methods

### Sample Collection

Walleye anal fin clips from Lake Winnipeg individuals were collected by boat electrofishing in Spring 2017 and 2018 from spawning sites in the Red River and possible spawning sites near Matheson Island and in the Dauphin River, representing the south basin, channel, and north basin, respectively (*n* = 129, 51, and 55 respectively). Prior to sampling, a Portable Electroanesthesia System (Smith Root) was used to anesthetize fish following approved animal use protocols of Fisheries and Oceans Canada (FWI-ACC-2017-001, FWI-ACC-2018-001), the University of Manitoba (F2018-019), and the University of Nebraska-Lincoln (Project ID: 1208). Fin clips were stored in 95% ethanol. For Lake Manitoba fish, *n* = 50 fry from the 2019-year class were sampled at Swan Creek Hatchery and stored in 95% ethanol. From Cedar Bluff Reservoir, anal fin clips were taken from *n* = 60 sexually mature walleye (*n* = 40 females and *n* = 20 males). Fin clips were collected after gametes were harvested. Procedures for sample collection at Cedar Bluff Reservoir were approved by the Fort Hays State University Institutional Animal Care and Use Committee (Protocol #19-0023). Fish were captured using seven 25 mm-mesh trap nets and four 76 mm mesh gill nets (91.44 X 1.83 m panel per net). In total, *n* = 345 walleye were genotyped for the present study. See Figure 1 for information on sampling locations.

### DNA Extraction, Sequencing, and Single Nucleotide Polymorphism Calling

DNA was extracted using the Qiagen DNeasy Blood & Tissue Kit (QIAGEN, Venlo, Netherlands) following manufacturer protocols. RAD-seq library preparation was performed with the NEBNext Ultra DNA Library Prep Kit for Illumina (New England Biolabs, Ipswich, Massachusetts, USA) with the *PstI* restriction enzyme. BestRAD libraries were baited using protocols outlined in Euclide et al., (2021) using a RAD-Capture panel for a different walleye research project in the Laurentian Great Lakes that was still ongoing at the time of publication (Ali et al., 2016). The panel includes a single bait for 99,636 SNP loci identified from a preliminary PSTI RAD-sequencing survey of 48 walleye collected in the Great Lakes (*n* = 9 – 10 from each of Lakes Superior, Huron, Michigan, Erie, and Ontario; USA) designed by Arbor Biosciences (Ann Arbor, Michigan, USA). Target loci were selected based on minor allele frequency (>0.01) and alignment position along a draft walleye genome to retain one SNP approximately every 5000 bp along every contig greater than 0.1 megabases. Bait-captured libraries were then sequenced on a S4 NovaSeq 6000 (Illumina, San Diego, California, USA) at NovoGene (Sacramento, California, USA).

Raw reads were processed with STACKS v2.3 (Catchen et al., 2013; Catchen et al., 2011), and aligned to the yellow perch (*Perca flavescens*) genome with bwa v0.7.17 (Feron et al., 2020; Li & Durbin, 2009). Single nucleotide polymorphisms (SNPs) were called following best practices with RAD data (Linck & Battey, 2019), and filtered for paralogy with HDPlot (McKinney et al., 2017). Data for population structure and demographic history were pruned for linkage with PLINK v1.9 (Purcell et al., 2007). Because relatedness can introduce bias in association studies, pairwise relatedness was checked using the method of moments as implemented by PLINK in SNPRelate v1.22 (Zheng et al., 2012). Because XP-EHH requires complete data, imputation of a set of SNPs with 10% missing data was performed with Beagle v5.1 (Browning et al., 2018; Browning & Browning, 2007; Weng et al., 2013; Yang et al., 2014). VCFtools v0.1.16, PGDSpider v2.1.5, the statistical computing environment R v4.0.3, along with the packages Tidyverse v1.3.0 and vcfR v1.12.0, were used throughout these analyses (Danecek et al., 2011; Knaus & Grünwald, 2017; Lischer & Excoffier, 2012; R Core Team, 2021; Wickham et al., 2019). Full details of SNP calling and processing are provided in the supplemental materials.

To further investigate predicted synteny between the walleye and yellow perch genomes, synteny was analyzed using progressiveMauve and i-ADHoRe following a pipeline published in Doerr & Moret (2018), with details provided in the supplementary materials (yellow perch genome, PLFA_1.0 at NCBI SRA BioProject PRJNA514308; walleye genome, ASM919308v1 at NCBI SRA BioProject PRJNA528354) (Darling et al., 2010; Ghiurcuta & Moret, 2014; Krzywinski et al., 2009; Proost et al., 2012).

### Population Genetics

Admixture v1.3.0 was run over K values of one through six, with 10-fold cross validation and 1000 bootstrap replicates for parameter standard errors (Alexander et al., 2009). Pophelper v2.3.0 in R was used to visualize and organize Admixture results (Francis, 2017). For population assignments, individuals were considered assigned to a cluster when their Q-values were > 0.85 for that cluster. Only assigned individuals were used for population differentiation and demographic reconstruction.

Hierfstat v0.5-7 was used to find β_WT_ as a measure of population-specific differentiation from the entire pool and Weir & Cockerham’s pairwise *F*_ST_ between populations (Goudet, 2005; Weir & Cockerham, 1984; Weir & Goudet, 2017). Ninety five percent confidence intervals were generated for *F*_ST_ and β_WT_ over 1,000 bootstrapped iterations each. In addition, observed heterozygosity (H_O_), gene diversity (H_S_), and inbreeding coefficients (*F*_IS_, 95% confidence intervals generated over 1,000 bootstrapped iterations) for each population were calculated.

NeEstimator v2.1 was used to estimate effective population size in 95% confidence intervals for each population using the linkage disequilibrium method and only comparing SNPs on different chromosomes (Do et al., 2014). Here, datasets were filtered for 10% missing data per population using population assignments with vcftools, pulled from the pruned SNP dataset.

EASYPOP v2.0.1 (Balloux, 2001) was used to test different scenarios of historic gene flow that may have led to contemporary *F*_ST_ between Lake Winnipeg and Lake Manitoba. Migration rates varied between 0, 0.0001, 0.001, 0.002, 0.003, 0.004, 0.005, 0.01, 0.02, and 0.03 individuals per generation, with migration rates held constant within each simulation run.

Population sizes of 4,500 individuals in Lake Winnipeg and 350 individuals in Lake Manitoba were assumed constant with equal males and females, based on estimates of N_e_ (see results). 1,000 simulated loci with two allelic states were used with free recombination allowed, using a KAM mutation model with mutation rate µ of 3.28x10^-9^ based on divergence between the blackfin icefish (*Chaenocephalus aceratus*) and dragonfish (*Parachaenichthys charcoti*) (Kim et al., 2019). Maximal variability (i.e., randomly assigned alleles) was chosen for the initial population. Each simulation was run over 930 generations, based on the 4.3 year generation time estimated in Franckowiak et al. (2009) and an approximate lower bound of 4,000 radiocarbon years since Lake Winnipeg had a reduced northern lakebed (Lewis et al., 2002). This reduced northern lakebed would have prevented walleye migration between populations via the Dauphin River and Lake St. Martin in the north basin (Fig 1). 100 replicate runs of EASYPOP were used for each migration rate tested, with 1000 simulation runs total. For each run, Weir & Cockerham’s pairwise *F*_ST_ was measured with Hierfsat. These *F*_ST_ values were compared to the *F*_ST_ observed between Lake Winnipeg and Lake Manitoba walleye populations using linkage disequilibrium-pruned SNPs.

### Signatures of Selection

Signatures of selection, where one genomic region has approached or reached fixation in one population compared to another, were explored with XP-EHH using the R package rehh v3.2.1, with the markers previously described as phased and imputed after filtering for 90% present data (Gautier et al., 2017; Sabeti et al., 2007). Data were unpolarized for analyzing XP-EHH because ancestral and derived alleles were unknown. Individuals with phased and imputed SNPs were split into the three assigned populations using Admixture results (Q>0.85 per individual), and extended haplotype homozygosity (EHH) was analyzed for each population separately (scan_hh). Three pairwise comparisons between each population were used to identify XP-EHH using false discovery rate corrected *p*-values (*q*-values). Significant candidate regions of selection were identified by analyzing 100 kilobase (kb) windows overlapping by 10 kb, in which at least three significant SNPs (*q* < 0.05) showing XP-EHH were found. For visualization, only significant SNPs within candidate regions were highlighted (thus a significant SNP outside a candidate region would be de-emphasized), using the R package ggman v0.99.0 with relative SNP positions to reflect distances between points in the reduced representation data (https://rdrr.io/github/veera-dr/ggman/). Results for XP-EHH were divided into three pairwise tests between each assigned population (Cedar Bluff Reservoir, Lake Manitoba, and Lake Winnipeg), and significant results for each pairwise test were further categorized into which population showed elevated XP-EHH scores (e.g., in a pairwise test between Lake Manitoba and Lake Winnipeg, certain SNPs were significant among haplotypes in Lake Manitoba versus others significant among haplotypes in Lake Winnipeg).

The program pcadapt v4.3.3 was used to explore potential signatures of local adaptation with outlier tests (Luu et al., 2017; Privé et al., 2020). Here, a scree plot showed that the optimal choice for principal components (PC) was K = 2 based on Cattell’s rule, an observation confirmed by a lack of discernible populations structure in K > 2 principal components (Cattell, 1966) (Fig S1). Significance for individual SNPs was determined using *q* values implemented in the R package qvalue v2.20.0, where a SNP was accepted as significant at *q* < 0.05 (Storey et al., 2020). The PC associated with a significant SNP was retrieved with *get.pc*, which enabled us to link significant SNPs with different axes of population structure. pcadapt results were separated into datasets of those significant along PC1 or PC2, corresponding to latitude and longitude, respectively.

Hierfstat v0.5-7 was used to provide supporting evidence for selection at individual SNPs using *F*’_ST_ with *basic.stats* on the same set of SNPs used with pcadapt. Here, *F*’_ST_ was used as a sample size corrected *F*_ST_ and measured in pairwise comparisons between each of three populations defined by admixture. In addition, absolute allele frequency differences were analyzed between populations. VCFtools was used to calculate allele frequencies for each population. Minor alleles in Lake Winnipeg-assigned individuals were used for comparisons across populations.

### Putative Inversion Analysis

To assess evidence that a region of interest that we identified (chromosome 8, position 15,260,000 - 15,900,000; see results for more detail) was a putative inversion, we conducted the following analyses as suggested by Huang et al. (2020) and Shi et al. (2021). First, to test whether the region showed elevated linkage disequilibrium or LD (*r*^2^), we calculated *r*^2^ using PLINK v1.9 for SNPs on chromosome 8 (--r2 --ld-window-r2 0.05 --ld-window 999999999 --ld-window-kb 30000) and generated a LD heatmap (Chang et al., 2015; Purcell et al., 2007). Second, we conducted regional PCA using SNPs within the region to identify whether individuals were grouped into three clusters along PC1, which is characteristic of chromosomal inversions. The discreteness of the clustering was calculated as the proportion of the between-cluster sum of squares over the total using the R function *kmeans* in *adegenet* (Jombart, 2008). Third, we compared heterozygosity (the proportion of heterozygotes) among the three identified PCA clusters using Wilcoxon tests (α = 0.05) to further confirm that individuals in the middle PCA cluster was heterozygous for the putative inversion. Lastly, we calculated genotype frequencies of the putative inversion in all populations and assumed that the more derived inversion arrangement would have had lower heterozygosity given its relatively recent origin compared to the ancestral state (Knief et al., 2016; Laayouni et al., 2003; Twyford & Friedman, 2015). Individual assignments for the three identified PCA clusters for the putative inversion were then visualized with respect to population assignments and site collected, to assess the frequency of the inversion across the different waterbodies studied.

To provide evidence of potential functional significance for the identified putative inversion on chromosome 8, gene expression data was collected for genes within the inversion. RNA-seq data was used from *n*=48 walleye in three sites in Lake Winnipeg, using previously published reads (NCBI SRA database accession #PRJNA596986) (Thorstensen et al., 2020). Three genes in the yellow perch genome were within the putative inversion on chromosome 8: *PDHX*, *EHF*, and *LRRC4C*. Therefore, expression of these genes was assessed with counts per million (CPM) values from the walleye transcriptome-aligned data (Jeffrey et al., 2020; Patro et al., 2017; Soneson et al., 2015). Differential gene expression was not tested for because the putative inversion was nearly fixed in the Lake Winnipeg-assigned walleye; differential expression would thus be expected between populations and not within populations. Details for RNA-seq are provided in supplementary materials.

## Results

### DNA Extraction, Sequencing, and SNP Calling

96,955 SNPs were called in STACKS, of which 63,882 were accepted as non-paralogous SNPs after using HDPlot. Pruning non-paralogous SNPs for linkage with PLINK left 46,342 SNPs for use for population structure and demographic reconstruction. The dataset of 63,882 SNPs was filtered for a maximum of 10% missing data and imputed and phased into a dataset of 18,601 SNPs for use with XP-EHH. Synteny between the walleye and yellow perch genomes was high, indicating a large degree of concordance between the two genomes (mean weighted synteny score = 0.999, 95% CI [0.997, 1.0]; mean relaxed synteny sore = 0.999, 95% CI [0.997, 1.0]) (Fig S2).

### Population Genetics

Admixture identified K = 2 populations based on the lowest cross validation error between K = 1 through 6, separating Canadian and Kansas populations of walleye (Fig S3). However, cross validation error was similar at K = 3 and population structure delineated the three different water bodies used in the present study; individuals were thus assigned to populations based on K = 3. Using ancestry coefficients Q > 0.85 for population assignments, 60 individuals were assigned to the Kansas population, 67 individuals to the Lake Manitoba population, 189 individuals to a Lake Winnipeg population, and 29 to no population. Notably, 27 out of the 29 unassigned individuals were found in Lake Winnipeg (the 2 remaining in Lake Manitoba), and of those, 18 were caught in the northern Dauphin River, 6 in the central Matheson Channel, and 2 in the southern Red River. In addition, out of the 67 individuals assigned to the Lake Manitoba population, 20 were sampled from the Dauphin River in Lake Winnipeg.

Pairwise *F*_ST_ showed the greatest differentiation between each Canadian lake and Cedar Bluff Reservoir, and moderate differentiation between Lake Manitoba and Lake Winnipeg (Fig 2). Population-specific differentiation from the overall pool, β_WT_, was lowest for Cedar Bluff Reservoir (Fig S4). Filtering for 90% present data for NeEstimator v2 from the pruned SNP dataset left 11,784 SNPs available in with the Cedar Bluff Reservoir population, 16,436 SNPs in the Lake Manitoba population, and 13,213 SNPs in the Lake Winnipeg population. Linkage disequilibrium-based N_e_ was highest for the Lake Winnipeg population, and approximately 20-35x lower in each the Lake Manitoba and Cedar Bluff Reservoir populations (Lake Winnipeg N_e_ = 12,084, 95% CI [9,986, 15,282]; Lake Manitoba N_e_=336, 95% CI [331, 342]; Cedar Bluff Reservoir N_e_ = 587, 95% CI [568, 608]) (Fig 2). Observed heterozygosity, gene diversity, and inbreeding coefficient (H_O_, H_S_, and *F*_IS_, respectively) were each highest for the Cedar Bluff Reservoir population, with *F*_IS_ approximately two times higher for Cedar Bluff walleye than the walleye in the northern lakes (Fig 2, Fig S4).

**Figure 2.**
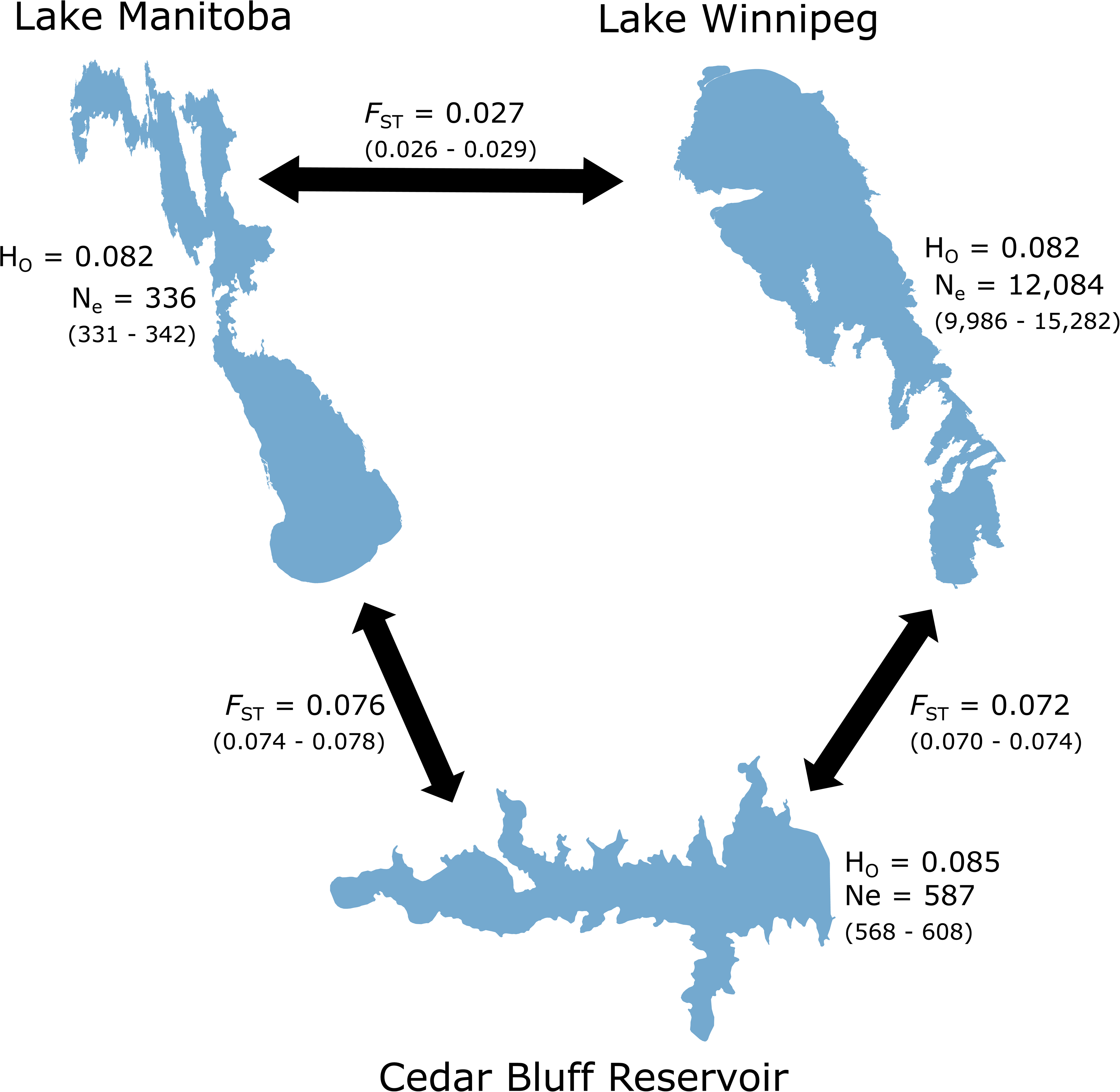
Population differentiation, effective population size, and observed heterozygosity for walleye (*Sander vitreus*) in three North American waterbodies. Cedar Bluff Reservoir (Kansas, USA) represents an entirely stocked population of walleye, while Lake Manitoba and Lake Winnipeg (Manitoba, Canada) walleye represent populations with possible gene flow. *F*_ST_ represents Weir & Cockerham’s pairwise *F*_ST_, while H_O_ observed heterozygosity, each found using Hierfstat. Effective population size for each population is represented by N_e_, found using NeEstimator v2. For *F*_ST_ and N_e_, 95% confidence intervals are provided in parentheses.

Simulations with EASYPOP showed observed *F*_ST_ between Lake Manitoba and Lake Winnipeg (between 0.026 and 0.029, Fig 2) most consistent with continuous migration of 0.001 individuals per generation starting 930 generations prior to the present (Fig S5).

### Signatures of Selection

The XP-EHH scores between Lake Winnipeg- and Lake Manitoba-assigned individuals identified 45 SNPs with elevated XP-EHH scores (15 SNPs higher in Lake Winnipeg, 30 higher in Lake Manitoba), of which 33 SNPs were in candidate regions under selection (11 with higher XP-EHH scores in Lake Winnipeg, 22 with higher XP-EHH scores in Lake Manitoba) (Fig 3).

**Figure 3.**
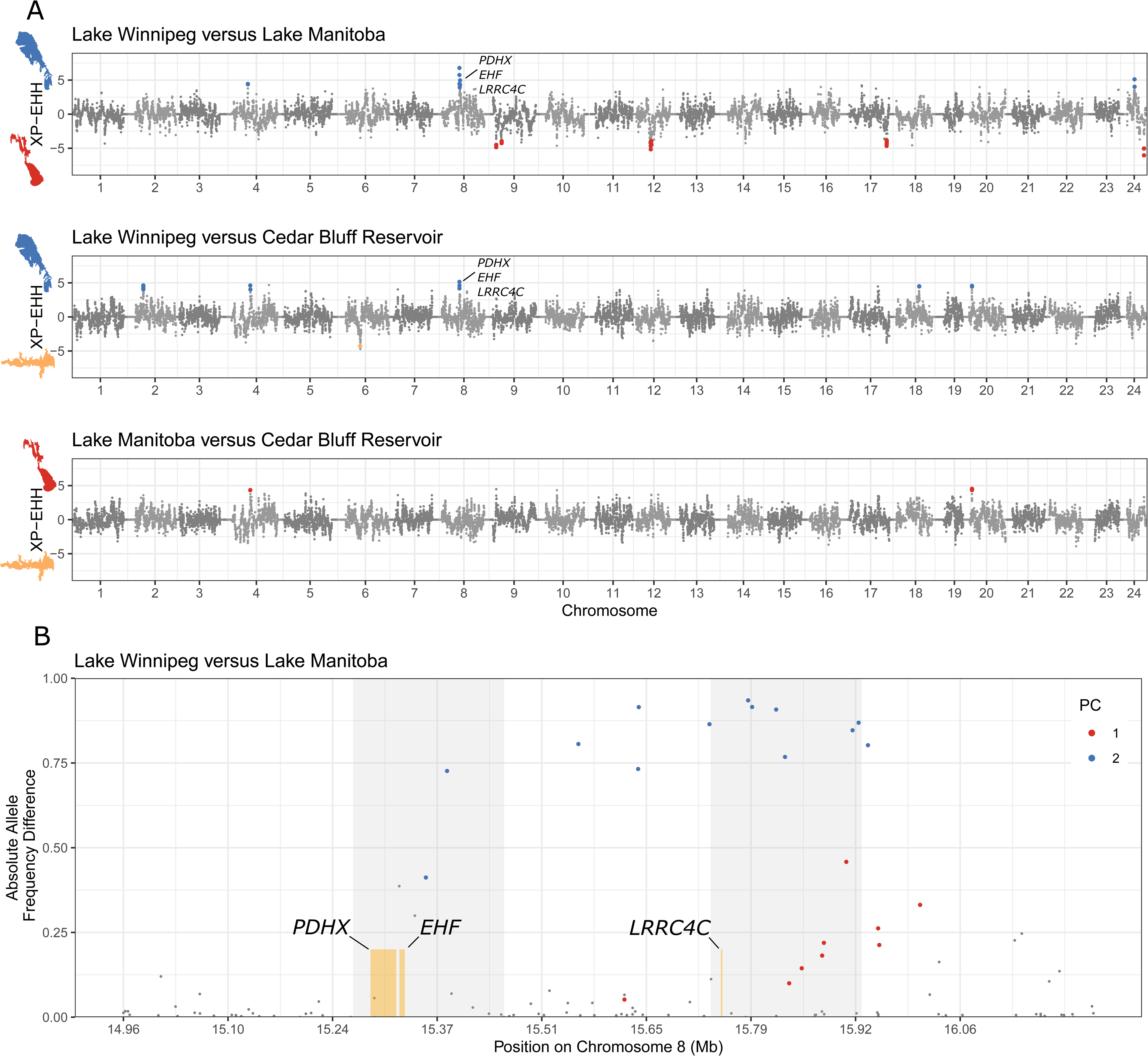
Signatures of selection inferred from cross-population extended haplotype homozygosity (XP-EHH) between each population of walleye (*Sander vitreus*) analyzed in the present study, along with absolute allele frequency differences between two waterbodies. In A), Cedar Bluff Reservoir (Kansas, USA) represents an entirely stocked population, while Lake Manitoba and Lake Winnipeg (Manitoba, Canada) each represent native populations with possible gene flow. Walleye DNA was aligned to the yellow perch (*Perca flavescens*) reference genome, and chromosome numbers represent yellow perch chromosomes. Unplaced scaffolds are not visualized. Single nucleotide polymorphisms (SNPs) were considered significant with *q*<0.05 and only if the SNPs were in a candidate region defined by 100 kilobase regions of the genome overlapping by 10 kilobases, in which ≥ 3 SNPs total were significant. Because XP-EHH focuses on haplotypes of unusual length and high frequency in one population compared to another, the population in which haplotypes are long and are at or approaching fixation can be identified. Here, the population in which XP-EHH was significant is identified with colours, where blue represents Lake Winnipeg, red represents Lake Manitoba, and yellow represents Cedar Bluff Reservoir. In B), absolute allele frequency differences were calculated between Lake Manitoba and Lake Winnipeg-assigned walleye. PC1 and PC2 refer to significant SNPs from principal component 1 or 2 from pcadapt (see Fig. 4). The genes *pyruvate dehydrogenase protein X component (PDHX), ETS homologous factor (EHF), and leucine-rich repeat-containing protein 4C (LRRC4C)* were within the candidate region for selection in Lake Winnipeg on chromosome 8. The candidate regions for selection from XP-EHH are emphasized with gray background, which together were identified as the putative chromosomal inversion.

The XP-EHH scores between Lake Winnipeg- and Cedar Bluff Reservoir-assigned individuals identified 26 SNPs with elevated XP-EHH scores (24 higher in Lake Winnipeg, 2 higher in Cedar Bluff Reservoir), of which 18 were in candidate regions under selection (17 with higher XP-EHH scores in Lake Winnipeg, 1 with higher XP-EHH scores in Cedar Bluff Reservoir) (Fig 3). Between Lake Manitoba- and Cedar Bluff Reservoir-assigned individuals, 7 SNPs showed elevated XP-EHH scores (7 with higher XP-EHH scores in Lake Manitoba, 10 with higher XP-EHH scores in Cedar Bluff Reservoir), of which 5 were in candidate regions under selection (5 with higher XP-EHH scores in Lake Manitoba, 0 with higher XP-EHH scores in Cedar Bluff Reservoir) (Fig 3). There were 10 candidate regions under selection in XP-EHH between Lake Winnipeg and Lake Manitoba, 5 candidate regions between Lake Winnipeg and Cedar Bluff Reservoir, and 0 candidate regions between Lake Manitoba and Cedar Bluff Reservoir (Table S1). Prominent among candidate regions was a section of chromosome 8 between approximately 15.26 Mb and 15.90 Mb in Lake Winnipeg with high differentiation compared to both Lake Manitoba and Cedar Bluff Reservoir.

Two principal components (PCs) showed population structure in pcadapt, with PC1 showing 5.76% variance explained by latitudinal differences between sites and PC2 showing 2.28% variance explained by longitudinal differences (Fig 4). 345 SNPs were identified as significant outliers by pcadapt, of which 226 were significant along PC1 and 119 were significant along PC2 (Fig 4). Along PC2, between approximately 15.35 Mb and 15.91 Mb, chromosome 8 showed a region of strong differentiation between Lake Manitoba and Lake Winnipeg.

**Figure 4.**
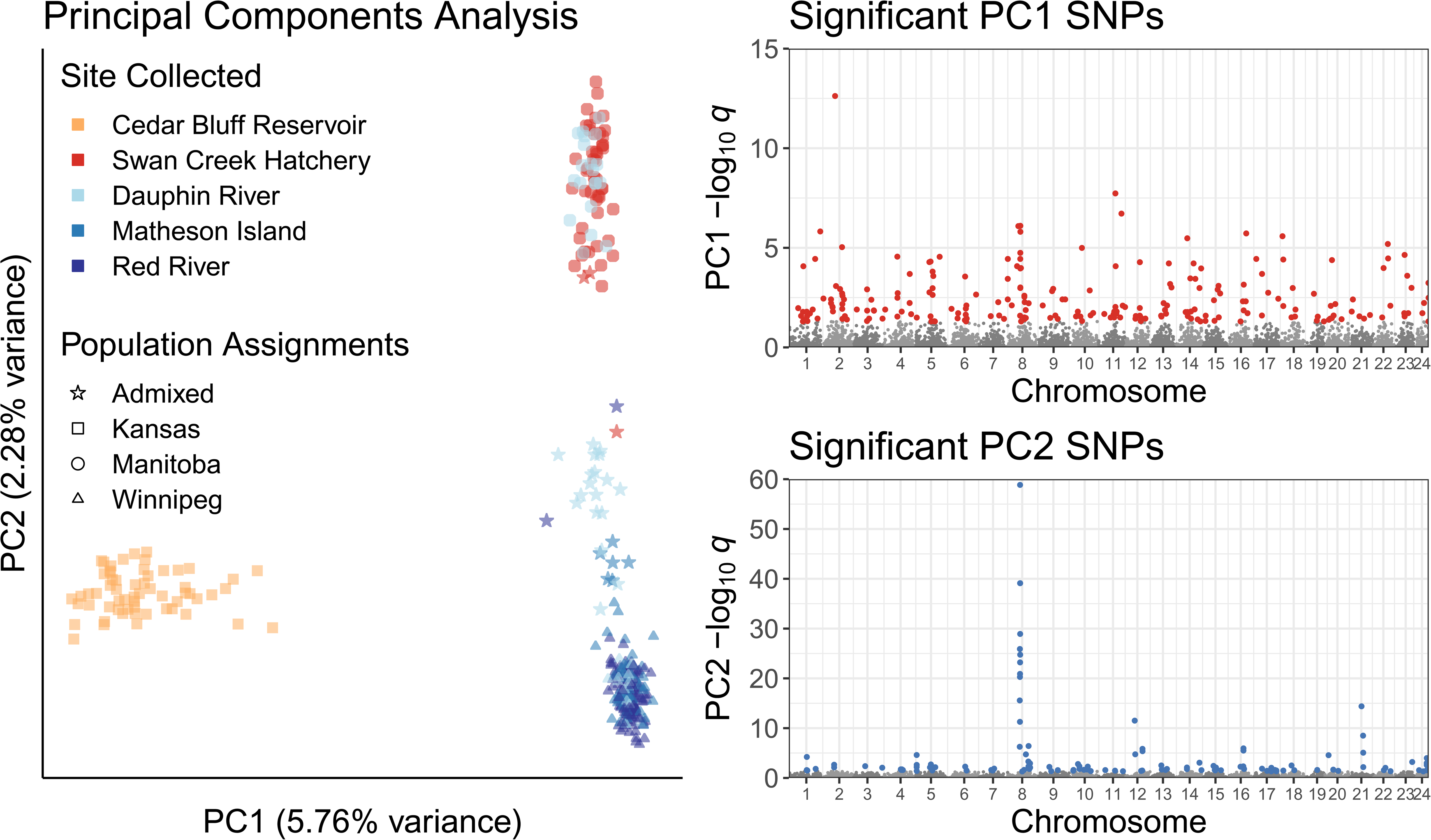
Principal components analysis of walleye (*Sander vitreus*) from three waterbodies across North America. Cedar Bluff Reservoir (Kansas, USA) represent an entirely stocked population of walleye of unknown origin. Lake Winnipeg and Lake Manitoba (Manitoba, Canada) represent native populations of walleye with possible gene flow. Principal components analysis was done using pcadapt, while population assignments were done using Admixture with K=3 groups. Individuals were assigned to a population with Q > 0.85 for a population, and individuals with maximum Q ≤ 0.85 were considered unassigned. Significant SNPs were distinguished according to the first two principal components in which they most strongly contributed variation, where principal component 1 (PC1) corresponded to latitudinal differences among populations, while principal component 2 (PC2) was most strongly characterized by variation between Lake Winnipeg and Cedar Bluff Reservoir relative to Lake Manitoba. Only SNPs with *q*-values < 0.05 were highlighted for visualization and retained for downstream analyses. Walleye DNA was aligned to the yellow perch (*Perca flavescens*) reference genome. Unplaced scaffolds were not visualized, and chromosome numbers refer to chromosomes in the yellow perch genome.

The region between 15.26 and 15.90 Mb along chromosome 8 was notable for its candidate regions identified in XP-EHH that were unique to Lake Winnipeg (Table S1), outlier SNPs in pcadapt (Fig 4), *F*^’^_ST_ outlier SNPs (Fig S6), and SNPs with absolute allele frequency differences (Fig 3, Fig S7) between Lake Manitoba and Lake Winnipeg. Many of the outlier SNPs in the chromosome 8 peak for *F’*_ST_ and absolute allele frequency differences overlapped with PC2 outlier SNPs from pcadapt at *q* < 0.05 (Fig 3, Fig 4, Fig S6, Fig S7).

### Putative Inversion Analysis

We found strong evidence of the genomic region 15.26 - 15.90 Mb on chromosome 8 as a putative chromosomal inversion (Fig 5). This region demonstrated elevated LD with a mean *r*^2^ of 0.30 and values up to 0.96 (Fig 5B). Regional PCA showed that individuals were clustered into three distinct groups along PC1 with a discreteness of 0.9626 (Fig 5C) and the middle PCA cluster displaying significantly higher heterozygosity than the other two clusters (Fig 5D, Fig S8). There were also significant differences in heterozygosity between the two homokaryotypes (Fig 5D) and the arrangement with lower heterozygosity (cluster 2) was assumed to be the derived inverted type. This putative inversion (cluster 2) was nearly fixed in the Red River and Matheson Island sites of Lake Winnipeg (toward the channel and south basin; Fig 1), was at intermediate frequency in the Dauphin River (north basin) of Lake Winnipeg, and was nearly fixed or heterozygous for the opposite genotype near Swan Creek Hatchery in Lake Manitoba and in Cedar Bluff (Fig 5E). Frequency of the putative inversion visualized in relation to individual assignments and site collected revealed the inversion was nearly fixed in Lake Winnipeg-assigned walleye for one genotype, nearly fixed for the other genotype among Lake Manitoba-assigned walleye, and among unassigned, likely admixed fish in Lake Winnipeg, nearly fixed for the Lake Winnipeg genotype (Fig S9).

**Figure 5.**
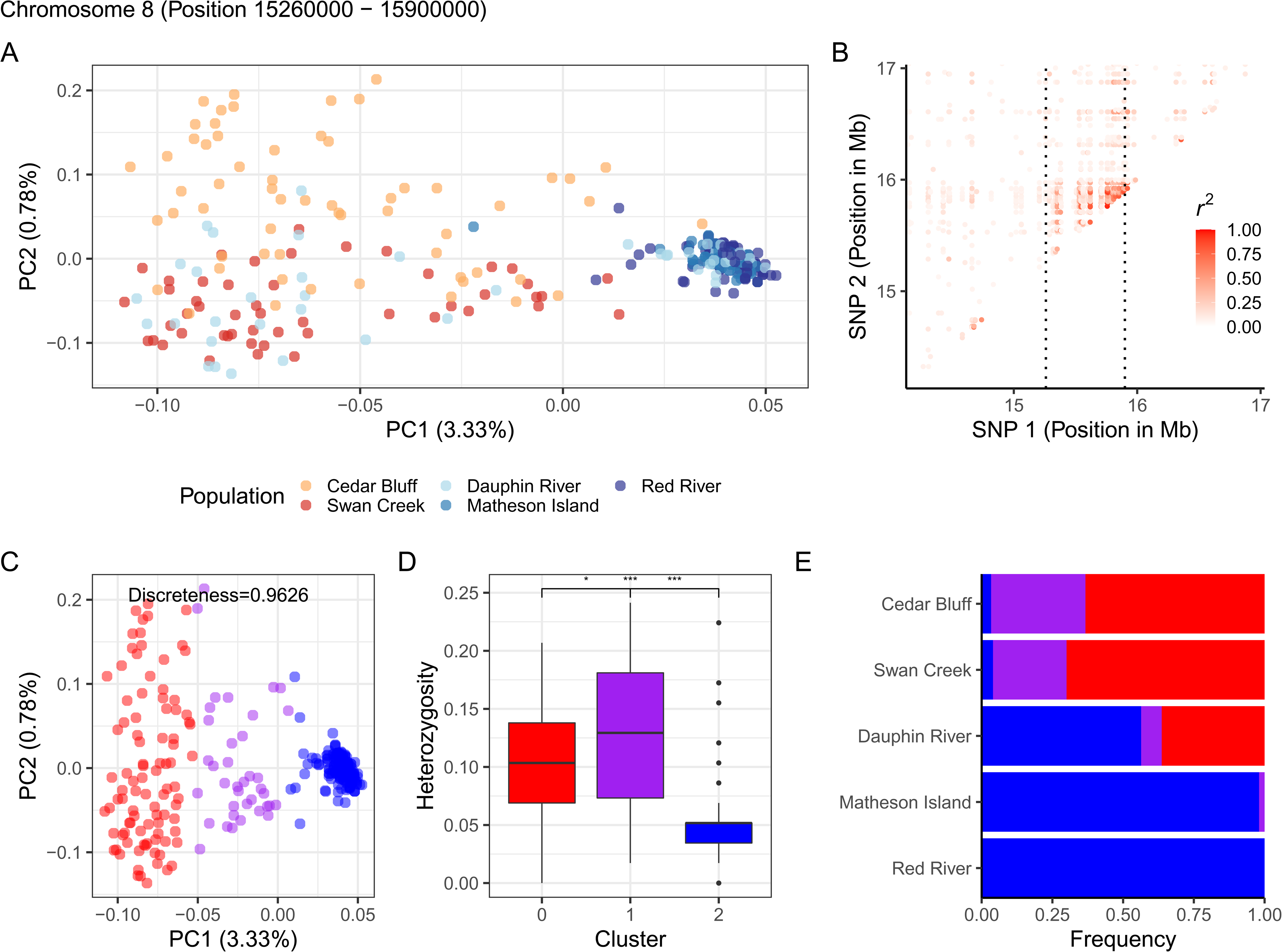
Characterization of the putative inversion on chromosome 8 (15.26 - 15.90 Mb). A) PCA based on SNPs within the putative inversion, coloured by site. B) Linkage disequilibrium heatmap for SNPs within the putative inversion, with boundary of the putative inversion emphasized by dashed lines. C) Clusters identified using k-means clustering represent two homozygous groups, red and blue, and a heterozygous purple group. D) Observed individual heterozygosity within each cluster identified. E) Genotype frequency distribution by collection site and cluster. Bars represent the proportion of individuals from each k-means cluster.

Three genes were identified within the putative inversion: *pyruvate dehydrogenase protein X component* (*PDHX*), *ETS homologous factor* (*EHF*), and *leucine-rich repeat-containing protein 4C* (*LRRC4C*). Each of these genes had transcripts expressed in Lake Winnipeg walleye, with counts per million between 0.4 and 20 (Fig S10).

## Discussion

We used a reduced representation approach to genotype *n* = 345 walleye from three different waterbodies in North America at 46,342 genetic markers. Walleye of the Canadian lakes diverged from each other despite three scenarios for gene flow: a connecting river, stocking of fry throughout the 20^th^ century, and flooding in 1882, 1902, 1904, 2011, and 2014 (and possibly periodically over evolutionary timescales). Historic gene flow likely occurred in sporadic pulses, as we observed evidence of limited but present contemporary gene flow possibly facilitated by flooding in 2011 and 2014. These potential pulses of gene flow were consistent with population differentiation (*F*_ST_) between the Canadian lakes that was lower in magnitude than each Canadian lake compared to a waterbody in Kansas, USA. Lake Winnipeg walleye showed unexpectedly strong signatures of selection on chromosome 8, with concurrent evidence for a chromosomal inversion between 15.26 and 15.90 Mb. An inversion may have played a role in adaptive divergence between Lake Winnipeg and Lake Manitoba walleye (Wellenreuther & Bernatchez, 2018; Shi et al., 2021), and three expressed genes (i.e., mRNA transcripts) within it indicate at least the potential for functional significance of the putative inversion.

### Population Genetics

Despite moderate signals of differentiation between the Canadian lakes, admixture analysis and population assignment identified 11% of individuals sampled from primarily the north basin of Lake Winnipeg as originating from Lake Manitoba. This gene flow into Lake Winnipeg may have been associated with stocking programs from Lake Manitoba to Lake Winnipeg in the 1900s and early 2000s, floods may have facilitated movement in 2011 and 2014 (Ahmari et al., 2016). However, individuals possibly admixed between Lake Manitoba and Lake Winnipeg were likely the direct descendants (i.e., F_1_ crosses) from individuals originating from each lake, based on admixture analyses (Bray et al., 2010; Reynolds & Fitzpatrick, 2013; Wilson & Goldstein, 2000). Therefore, we believe flooding in 2011 and 2014 is the more likely route for movement than stocking, because stocking ended in 2002, approximately 3.5 generations before sampling in 2017 and 2018, given an estimated 4.3 year generation time (Franckowiak et al., 2009). Simulations confirmed low gene flow between each lake, assuming continuous gene flow post-glaciation. Taken together, we suggest that gene flow between Lakes Manitoba and Winnipeg prior to stocking programs and recent flooding has been sporadic, and that several differentiated genomic regions suggestive of local adaptation remain in each population despite recent gene flow.

We observed moderate population differentiation (*F*_ST_) between two large lakes in Canada (Lake Winnipeg and Lake Manitoba), and comparatively stronger differentiation between the two lakes in Canada and Cedar Bluff Reservoir, Kansas, USA. This relatively stronger differentiation between the Canadian lakes used here and the waterbody in Kansas was similar in magnitude to differentiation observed between walleye in Minnesota and Wisconsin (USA), differences which were attributed to Pleistocene lineages (Bootsma et al., 2020). Walleye population differentiation between Lake Winnipeg and Lake Manitoba was consistent with certain other walleye populations within watersheds (e.g., Grand River, Ontario, Canada vs other Lake Erie stocks in Euclide et al., 2020; Sarah Lake vs Lake Koronis, Minnesota, USA & in Bootsma et al., 2020). Retreating glacial refugia dispersed walleye into different watersheds approximately 7,000 - 10,000 years ago (Bootsma et al., 2020; Stepien et al., 2009). Therefore, the population structure identified in the present study represents differentiation following geologically recent changes to the overall landscape. This differentiation is consistent with high spawning site fidelity previously observed in walleye (Horrall, 1981; Jennings et al., 1996; Stepien et al., 2009; Stepien & Faber, 1998), despite the species’ otherwise broad movements in their waterbodies (Munaweera Arachchilage et al., 2021; Raby et al., 2018; Turner et al., 2021).

Given widespread stocking and potamodromous movements in walleye, it may be surprising that the species showed as much population differentiation as has been observed between waterbodies in multiple watersheds (Bootsma et al., 2020; Euclide et al., 2021; Munaweera Arachchilage et al., 2021; Raby et al., 2018; Turner et al., 2021). While stocking has led to extensive homogenization in some walleye populations, the lack of gene flow from stocking in the present data and among other waterbodies implicates potential mechanisms by which gene flow is prevented (Bootsma et al., 2020). One possible mechanism is environmental. Here, environmental factors impede gene flow because local adaptation may select against migrants (Hendry, 2004; Sexton et al., 2014). In a comparison between environmental mechanisms of isolation and isolation by distance, isolation by environment was more common in all taxa, including vertebrates (Sexton et al., 2014). As such, environmental differences between source and stocked waterbodies may have precluded gene flow in different walleye systems. Genomic architecture is another possible mechanism that can maintain differentiation despite stocking. Here, physical linkage or chromosomal rearrangements have been hypothesized to play an important role in local adaptation in the face of gene flow (Tigano & Friesen, 2016). Walleye in waterbodies with historical pulses of gene flow or recent stocking may thus maintain differentiation because of genomic architecture facilitating local adaptation (Tigano & Friesen, 2016). Moreover, the hypotheses that environment or genomic architecture may have maintained differentiation despite opportunities for gene flow are not mutually exclusive. Because genomic architecture has played a role in local adaptation in several freshwater fishes (Shi et al., 2021), and environment is a key variable in local adaptation (Coop et al., 2010; Forester et al., 2018), these two mechanisms may often work in tandem.

### Putative Chromosomal Inversion

A putative chromosomal inversion was identified on chromosome 8. Based on PCA, absolute allele frequency differences, and *F*^’^_ST_, this 640 Kb region on chromosome 8 showed evidence as an outlier region between Lake Manitoba and Lake Winnipeg. For example, the top 10 outlier SNPs (by -log_10_ *q*-value) out of 119 significant outliers (*q* < 0.05) along PC 2 in a pcadapt analysis are all from this small outlier region (Luu et al., 2017; Privé et al., 2020). Similarly, the top 10 SNPs by absolute allele frequency difference between Lake Manitoba and Lake Winnipeg are also from this small outlier region. As a haplotype-based test, XP-EHH provided evidence that this region was selected for not in Lake Manitoba, but in Lake Winnipeg instead (Sabeti et al., 2007). This chromosome 8 region was also a consistent outlier with *F*_ST_ and BAYESCAN analyses (Foll & Gaggiotti, 2008) in a different study on walleye in Wisconsin and Minnesota (USA) (Bootsma et al., 2020). However, SNP data were too sparse to interrogate the genomic architecture of this outlier region. In the present study, a dense panel of SNPs provided sufficient resolution to investigate the possibility that this outlier region was a chromosomal inversion.

Linkage disequilibrium, regional PCA, discreteness of clustering, and heterozygosity provided evidence for a putative chromosomal inversion at higher frequency in Lake Winnipeg relative to both other waterbodies. This putative inversion was nearly fixed in Lake Winnipeg-assigned walleye, and at intermediate frequencies in walleye assigned to the other populations in this study. If the putative inversion had a neutral effect on walleye biology, an intermediate frequency would have been expected among admixed fish given the parental populations were almost fixed for opposite genotypes. Instead, among Lake Manitoba-assigned fish caught in Lake Winnipeg that we hypothesize migrated into Lake Winnipeg (the majority of which were found in the northern Dauphin River), the putative inversion was nearly fixed for the Lake Winnipeg genotype. Therefore, this putative inversion may play a role in walleye adaptation to Lake Winnipeg.

Three genes were within the putative inversion: *PDHX*, *EHF*, and *LRRC4C*. All three were expressed in Lake Winnipeg walleye, indicating at least some gene transcription activity within the putative inversion. However, the specific functional significance of these genes in Lake Winnipeg walleye is unknown. Several other characteristics of Lake Winnipeg walleye are consistent with signatures of selection unique to the waterbody. Observations of recently-evolved dwarf walleye morphotypes possibly due to fishing pressure suggest ongoing selection within Lake Winnipeg (Moles et al., 2010; Sheppard et al., 2018). Additionally, there are patterns of low gonadal investment despite high lipid concentrations in Lake Winnipeg walleye relative to walleye in other lakes, indicating a possibly unusual phenotype in walleye of Lake Winnipeg relative to others (Moles et al., 2008). Walleye habitats are different between Lake Manitoba and Lake Winnipeg, as well. Lake Winnipeg is one of the largest lakes in the world based on surface area, but is relatively shallow compared with other large freshwater lakes (Brunskill et al., 1980; ECCC & MARD, 2020). However, Lake Manitoba is shallower still, with a 4.9 m mean depth (compared with a mean depth for Lake Winnipeg of 13.3 m and 9 m in the north and south basins, respectively) (ECCC & MARD, 2020; Rawson, 1952). Drainage from the more saline Lake Manitoba increases salinity in Lake Winnipeg’s north basin, but not in Lake Winnipeg’s south basin (Brunskill et al., 1980; ECCC & MARD, 2020; ECCC, 2011). Consequently, possible local adaptation in Lake Winnipeg may be an evolutionary response to environmental differences in the waterbody.

## Conclusion

The identification of a putative inversion contributes to a broad literature on chromosomal inversions in diverse taxa (see Wellenreuther & Bernatchez, 2018), although information on inversions is relatively scarce for freshwater fishes (Shi et al., 2021). In addition, many observed inversions have been > 1 Mb in length, possibly because reduced representation sequencing methods may be biased for detecting large inversions (Wellenreuther & Bernatchez, 2018). The present data show a small putative inversion in an obligate freshwater fish discovered *via* reduced representation sequencing (i.e., Rapture). That this putative inversion may have played a role in recent divergence between fish of two connected lakes, and that its frequency among likely admixed fish is consistent with a possible adaptive role in Lake Winnipeg walleye, indicates a potential importance for chromosomal inversions in freshwater fishes more generally. Inversions have been previously related to environmental adaptation, such as in freshwater emerald shiner (*Notropis atherinoides*) with high gene flow (Wellenreuther & Bernatchez, 2018; Shi et al., 2021). Heterogeneous habitats with opportunities for gene flow are common for freshwater fishes (Griffiths, 2015; Mushet et al., 2019), and chromosomal inversions, along with other genomic architectures, may facilitate local adaptation in many organisms that live in such connected environments.

## Acknowledgements

We thank Dr. Jeff Long for contributing walleye fry from Swan Creek Hatchery and for valuable discussions of Manitoba walleye ecology, diversity, and stocking. We thank Colin Charles, Colin Kovachik, Doug Leroux, Nicole Turner, Mike Gaudry, Sarah Glowa, and Emily Barker, who provided fin clips used for walleye DNA from Lake Winnipeg. We thank Evelien de Greef for guidance on synteny analyses and the map figure, and Dr. Kristen Gruenthal for sharing her expertise in sequencing library preparation. We thank David Splausbury and his staff from the Kansas Department of Wildlife, Parks, and Tourism for their assistance in the collection of samples at Cedar Bluff Reservoir. We thank Dr. Colin Garroway for discussions about β_WT_. Many analyses in this manuscript were enabled using computing resources provided by WestGrid (www.westgrid.ca) and Compute Canada (www.computecanada.ca). This work was supported by a Fisheries and Oceans Canada Ocean and Freshwater Science Contribution Program Partnership Fund grant awarded to KMJ, JRT, and Dr. Darren Gillis, and a Natural Sciences and Engineering Research Council of Canada Discovery Grants awarded to KMJ (#05479) and JRT (#06052). Work by JRT is also supported by the Canada Research Chairs program (#223744) and the Faculty of Science, University of Manitoba (#319254).

## Data Accessibility

All metadata and scripts used for analyses in this manuscript are available at: https://github.com/BioMatt/walleyeDNA. Raw sequence reads are available at the National Center for Biotechnology Information Sequence Read Archive (accession #PRJNA800092).

## Author Contributions

Matt J. Thorstensen conducted the analyses and wrote the original manuscript. Matt J. Thorstensen, Jennifer D. Jeffrey, Ken M. Jeffries, Jason R. Treberg, Douglas A. Watkinson, and Eva C. Enders conceived of the study. Eva C. Enders, Douglas A. Watkinson, and Yasuhiro Kobayashi contributed to obtaining samples. Peter T. Euclide and Wesley A. Larson developed the sequencing panel used. Yue Shi analyzed the putative inversion. Ken M. Jeffries and Jason R. Treberg acquired funding. All authors participated in editing the manuscript and contributed to interpretation and analyses.

## Notes

https://github.com/BioMatt/walleyeDNA

### Competing Interest Statement

The authors have declared no competing interest.

### Summary of Updates

Grammar, figure captions, clarifications of text, and author contributions have been updated and in some cases corrected.

