## supplemental materials for "A chromosomal inversion may facilitate adaptation despite periodic gene flow in a freshwater fish"

### Supplemental Methods

*SNP Calling and Processing*

Raw reads were processed with STACKS v2.3 (Catchen et al., 2013; Catchen et al., 2011) using settings in Euclide et al., (2021), with the exception that the yellow perch (*Perca flavescens*) reference genome was used to align reads with bwa v0.7.17 with default settings (PLFA_1.0 at the National Center for Biotechnology Information (NCBI) Sequence Read Archive (SRA) BioProject PRJNA514308) (Feron et al., 2020; Li & Durbin, 2009) instead of the loci developed in Euclide et al., (2021). The closely-related yellow perch’s reference genome was used because we sought to link SNPs with annotated genes and an annotated walleye reference genome was not available at the time. In addition, the yellow perch reference genome was assembled at the chromosome-level, and the yellow perch and walleye share karyotypes (Bootsma et al., 2020; Danzmann, 1979). Loci were processed with *process_radtags* (-e pstI -i gzfastq -c -q -r --filter_illumina --bestrad -t 140), aligned with bwa, genotyped in *gstacks*, and SNPs were called with *populations* with one random SNP called per locus, a minor allele count ≥2, a ≥60% genotyping rate per site collected, and a ≥60% genotyping rate overall (--min-samples-per-pop 0.6 --min-samples-overall 0.6 --min-mac 2 --write-random-snp). As opposed to a higher minor allele frequency filter, singletons were excluded from these analyses following best practices for detecting population structure with reduced representation data (Linck & Battey, 2019). The SNPs were then filtered for paralogs using the heterozygosity and read ratio deviation method (HDPlot) where SNPs were accepted as non-paralogs with heterozygosity ≤0.60 and read ratio deviation ≤|7| (McKinney et al., 2017). For population structure and demographic reconstruction, the SNP dataset was pruned for linkage with PLINK v1.9 in 50 SNP windows shifted by 10 SNPs per step, filtered at an r^2^ threshold of 0.1 (--indep-pairwise 50 10 0.1) (Purcell et al., 2007). XP-EHH requires complete data, while phasing vastly increases statistical power in the approach (Gautier et al., 2017). Therefore, Beagle v5.1 was used to phase and impute unpruned SNPs because of its robust performance on a variety of datasets and observed 95% imputation accuracy with as low as 40% missing data (Browning et al., 2018; Browning & Browning, 2007; Weng et al., 2013; Yang et al., 2014). However, the overall SNP dataset was filtered to only 10% missing data for imputation and phasing. Because relatedness can introduce bias in association studies, pairwise relatedness was checked using the method of moments as implemented by PLINK in SNPRelate v1.22 (Zheng et al., 2012). Where appropriate, VCFtools v0.1.16 was used to filter or subset variant call format files, while PGDSpider v2.1.5 was used to convert data between formats (Danecek et al., 2011; Lischer & Excoffier, 2012). The statistical computing environment R v4.0.3, along with the packages Tidyverse v1.3.0 and vcfR v1.12.0, were used throughout these analyses (Knaus & Grünwald, 2017; R Core Team, 2021; Wickham et al., 2019).

#### Synteny in Walleye and Yellow Perch Genomes

Because SNP relationships with gene locations were important for functional analyses of signatures of selection and the yellow perch genome was used for these walleye data, synteny between available walleye and yellow perch genomes was investigated using a pipeline published in Doerr & Moret (2018) (yellow perch genome, PLFA_1.0 at NCBI SRA BioProject reference #PRJNA514308; walleye genome, ASM919308v1 at NCBI SRA BioProject reference #PRJNA528354) (Feron et al. 2020). First, progressiveMauve v20150226 build 10 was used to create a backbone to anchor genomic blocks using both genomes (Darling et al. 2010). Then i-ADHoRe v3.0.01 was used to analyze synteny in colinear mode with default settings (Proost et al. 2012). Circos v0.69-8 was then used to visualize synteny as an ideogram, while scripts published in Doerr & Moret (2018) were used to generate both relaxed and weighted synteny scores, along with being useful for data conversion between prior steps (scripts accessed from https://github.com/danydoerr/large_syn_workflow) (for Circos, see Krzywinski et al. 2008; for synteny scores, Ghiurcuta & Moret 2014).

*RNA Sequencing*

To quantify counts per million for genes within the putative inversion, *n*=48 female or sex-unidentified walleye were sampled sub-lethally for gill tissue at the Lake Winnipeg Red River, Matheson Island, and Dauphin River sites in May of 2017 and 2018 (*n*=8 fish per site per year; Fig 1) as in Thorstensen et al. (2020). Total RNA was extracted from gill samples preserved in RNA*later* (Thermo Fisher Scientific, Waltham, MA, USA) using the RNeasy Plus Mini Kit (Qiagen, Venlo, Netherlands) with minor modifications specified in Thorstensen et al. (2020).  Library preparation and sequencing was performed by Genome Québec (http://gqinnovationcenter.com), as in Thorstensen et al. (2020). Sequencing was performed using NovaSeq 6000 (Illumina) on a single lane and produced 2.17 billion paired-end reads, submitted to the NCBI SRA database (accession #PRJNA596986).

To quantify gene-level abundance, transcript abundance was quantified with Salmon v0.13.3 (Patro et al., 2017), using a previously assembled reference transcriptome for walleye generated by Jeffrey et al. (2020) (Sequence Read Archive Accession #SRP150633). The R/Bioconductor package “tximport” (Soneson et al., 2016) was used to estimate counts from abundances at the gene-level that were scaled using the average transcript length, averaged over samples, and library size (*countsFromAbundance = lengthScaledTPM*).

### Supplemental Results

| Comparison | Chromosome | Start | End | Number of Markers |
| --- | --- | --- | --- | --- |
| Lake Winnipeg v Lake Manitoba | 4 | 15,420,000 | 15,590,000 | 9 |
|  | **8** | **15,260,000** | **15,450,000** | **5** |
|  | **8** | **15,710,000** | **15,900,000** | **6** |
|  | 9 | 4,430,000 | 4,610,000 | 9 |
|  | 9 | 8,460,000 | 8,650,000 | 4 |
|  | 12 | 14,270,000 | 14,460,000 | 8 |
|  | 12 | 14,610,000 | 14,900,000 | 8 |
|  | 17 | 28,530,000 | 28,740,000 | 10 |
|  | 24 | 8,500,000 | 8,600,000 | 3 |
|  | 24 | 16,030,000 | 16,220,000 | 3 |
| Lake Winnipeg v Cedar Bluff Reservoir | 2 | 13,800,000 | 14,000,000 | 11 |
|  | 4 | 17,220,000 | 17,410,000 | 6 |
|  | 8 | 15,260,000 | 15,450,000 | 5 |
|  | 18 | 19,570,000 | 19,730,000 | 6 |
|  | 20 | 6,280,000 | 6,470,000 | 8 |

**Table S1** Significant candidate regions for cross-population extended haplotype homozygosity (XP-EHH). The comparison represents the two populations used for comparing walleye (*Sander vitreus*) populations. Cedar Bluff Reservoir (Kansas, USA) represents an entirely stocked population, while Lake Manitoba and Lake Winnipeg (Manitoba, Canada) represent native populations with possible gene flow. Candidate genomic regions for XP-EHH were identified by analyzing 100 kilobase (kb) windows overlapping by 10 kb, in which at least three significant SNPs (*q* < 0.05) showing XP-EHH were found. Chromosome represents the chromosome from the yellow perch (*Perca flavescens*) reference genome that walleye DNA was aligned to. Start and end represent beginning and ending base pair positions in the corresponding chromosome, defining the candidate region for selection. Number of markers shows the number of single nucleotide polymorphisms identified in the corresponding genomic region. In bold are two candidate regions identified as a putative chromosomal inversion.

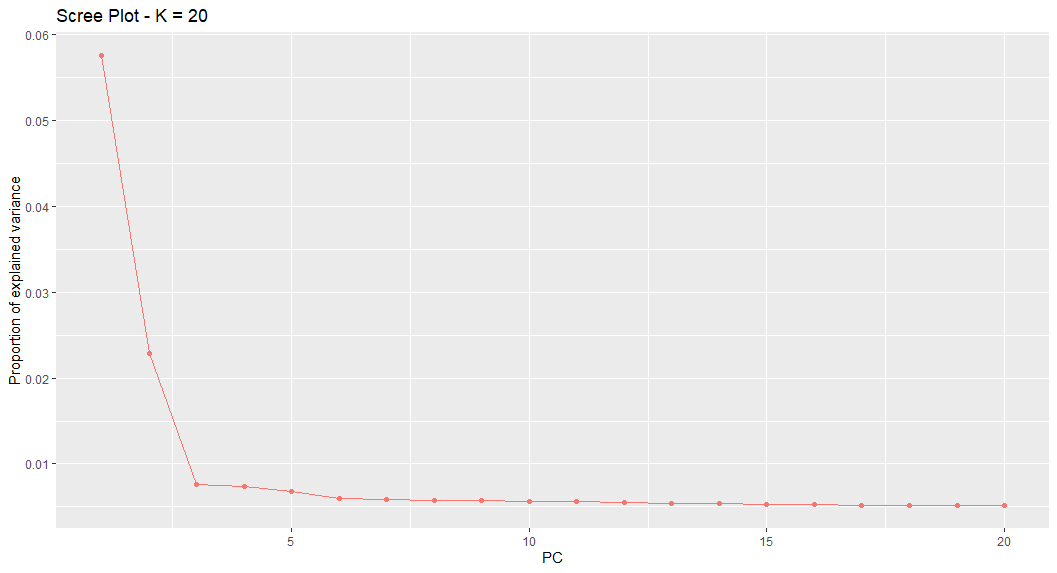

**Fig S1** Scree plot for proportion of explained variance in principal components analysis using pcadapt for walleye (*Sander vitreus*). By Cattell’s Rule, K=2 principal components explains a large proportion of variance in the data. These two principal components are used for subsequent analyses of outlier single nucleotide polymorphisms.

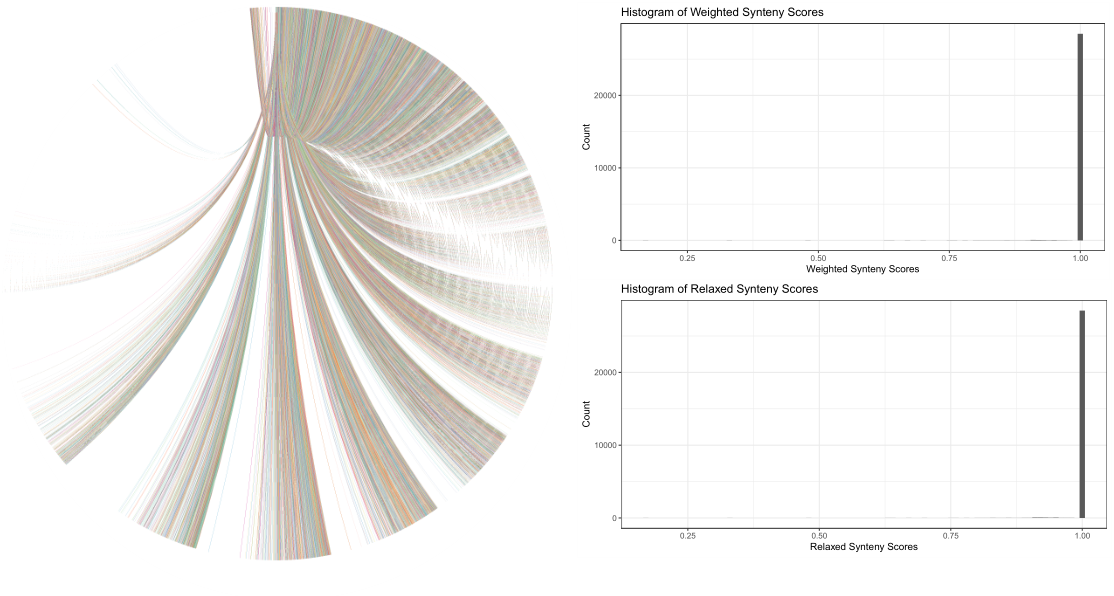

**Fig S2** Synteny analyses between the yellow perch (*Perca flavescens*) and walleye (*Sander vitreus*) genomes. The Circos plot is based on synteny analyses by the programs progressiveMauve and i-ADHoRe. Relaxed and weighted synteny scores were analyzed from those alignments, where higher values near 1 indicate greater complete synteny between the genomes.

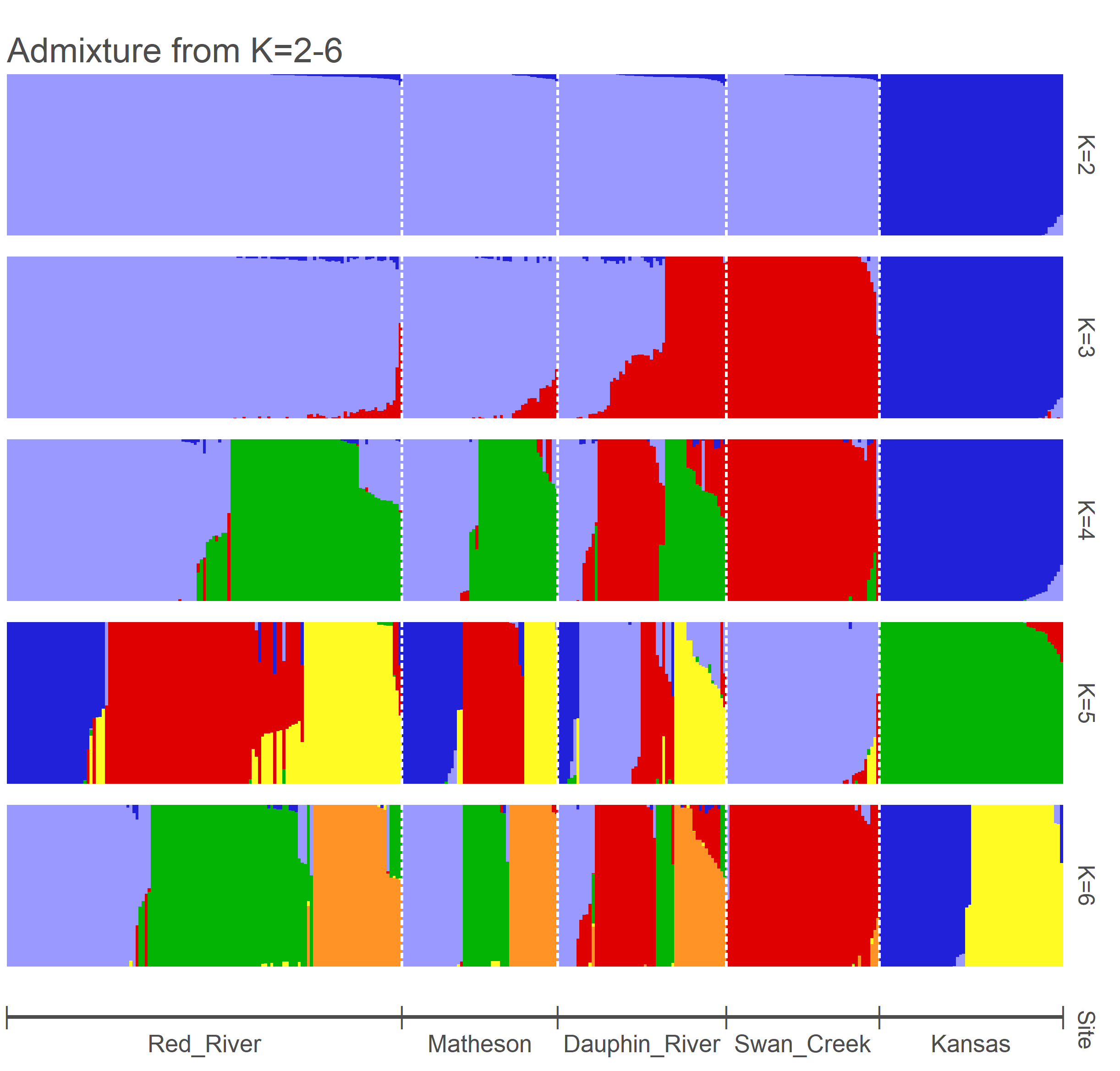

**Fig S3** Admixture results for *n*=345 walleye (*Sander vitreus*) sampled from Cedar Bluff Reservoir (Kansas, USA), Lake Manitoba (Manitoba, Canada), and Lake Winnipeg (Manitoba, Canada). The Red River, Matheson Island, and Dauphin River sites represent south basin, narrows, and north basin sites in Lake Winnipeg. Swan Creek Hatchery represents walleye from Lake Manitoba. Cedar Bluff Reservoir walleye are labeled as originating from Kansas in the present visualization. K=2 through 6 groups were tested for admixture analysis and population assignment.

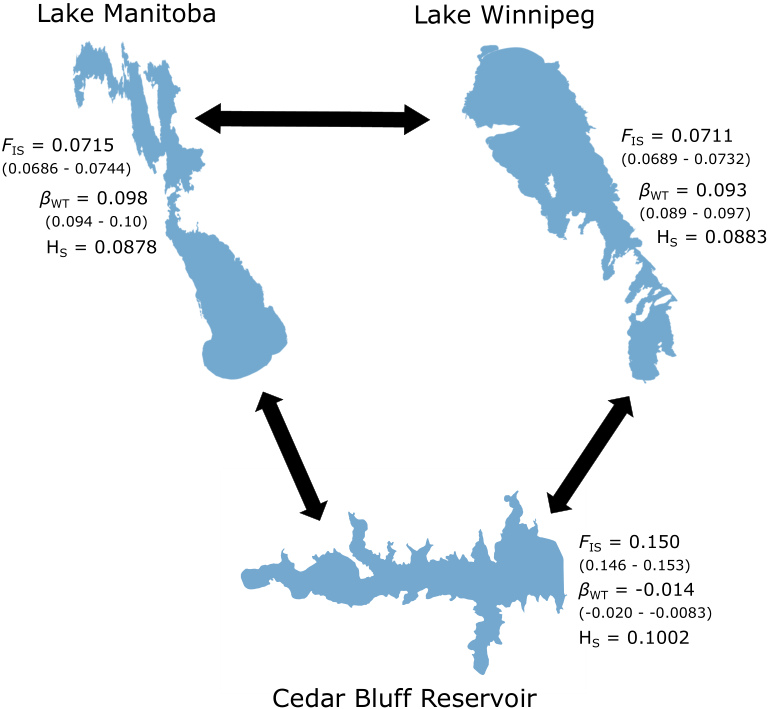

**Fig S4** Inbreeding coefficients (*F*_IS_), differentiation from the overall genetic pool (*β*_WT_), and gene diversity (H_S_) for three populations of walleye (*Sander vitreus*) from three waterbodies in North America. Cedar Bluff Reservoir (Kansas, USA) represents an entirely introduced walleye population of unknown origin, while Lake Manitoba and Lake Winnipeg (Manitoba, Canada) walleye are native and may experience gene flow between them. For *F*_IS_ and *β*_WT_, 95% confidence intervals are provided in parentheses.

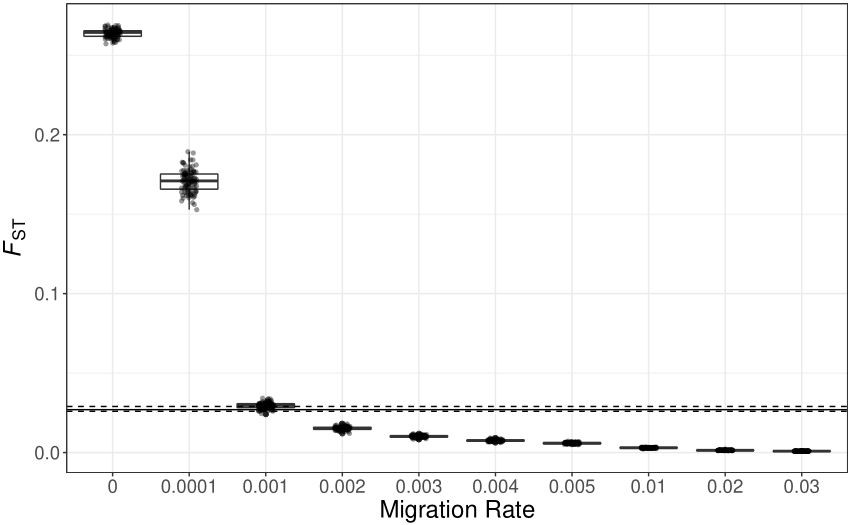

**Fig S5** EASYPOP simulated *F*_ST_ between Lake Manitoba and Lake Winnipeg populations of walleye_._ Migration rate represents simulated migration rates set in the program, while *F*_ST_ represents Weir & Cockerham’s pairwise *F*_ST_ among simulations. Each point represents one simulation run, with 100 replicate runs per migration rate. The solid horizontal line represents estimated *F*_ST_ from the data, while the dashed lines represent 95% confidence intervals from estimated *F*_ST_ (estimated *F*_ST_ = 0.027, 95% CI [0.026, 0.029]; Fig 2).

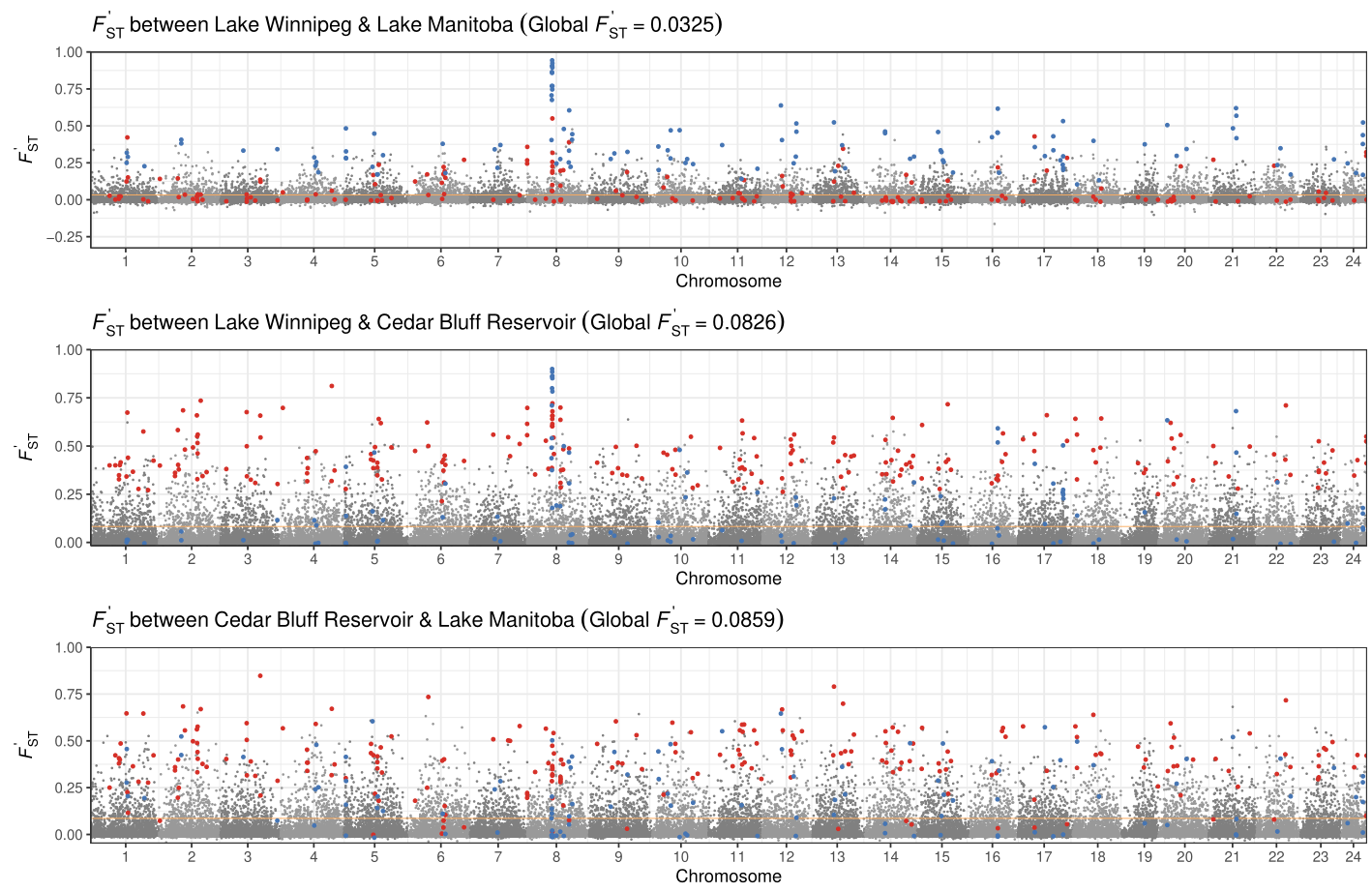

**Fig S6** *F’*_ST_ between assigned three assigned populations.

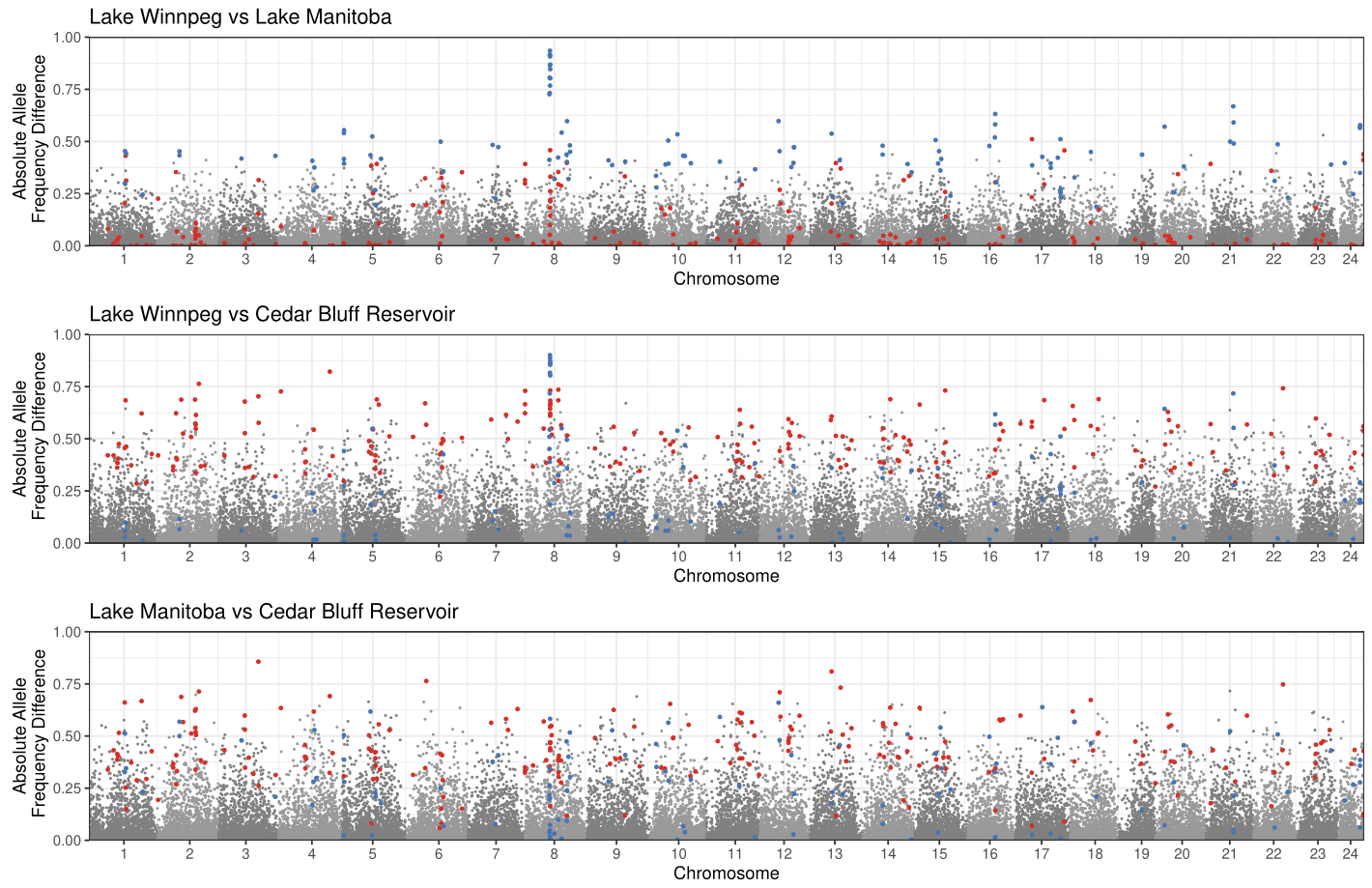

**Fig S7** Absolute allele frequency differences between three assigned populations.

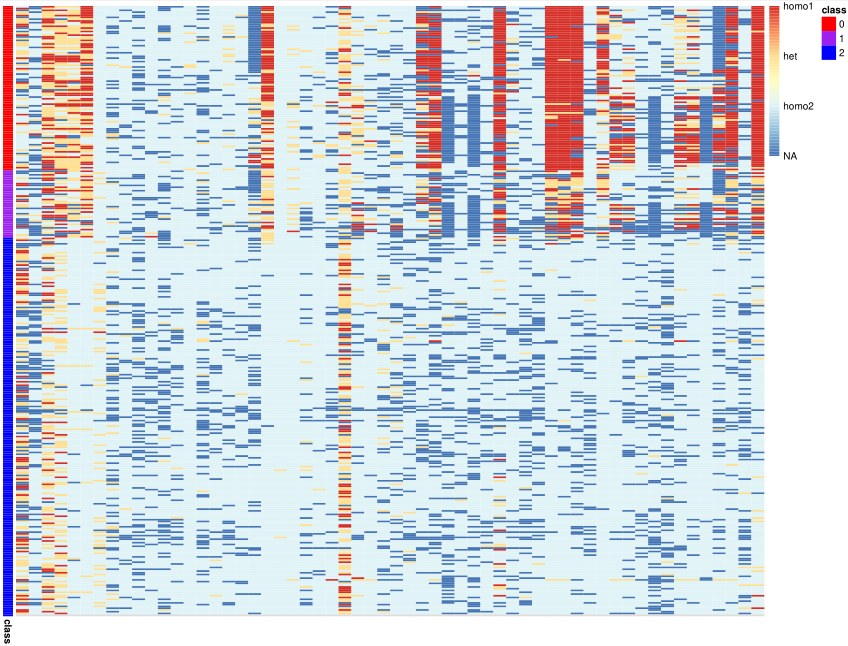

**Fig S8** Genotype heatmap within the putative inversion.

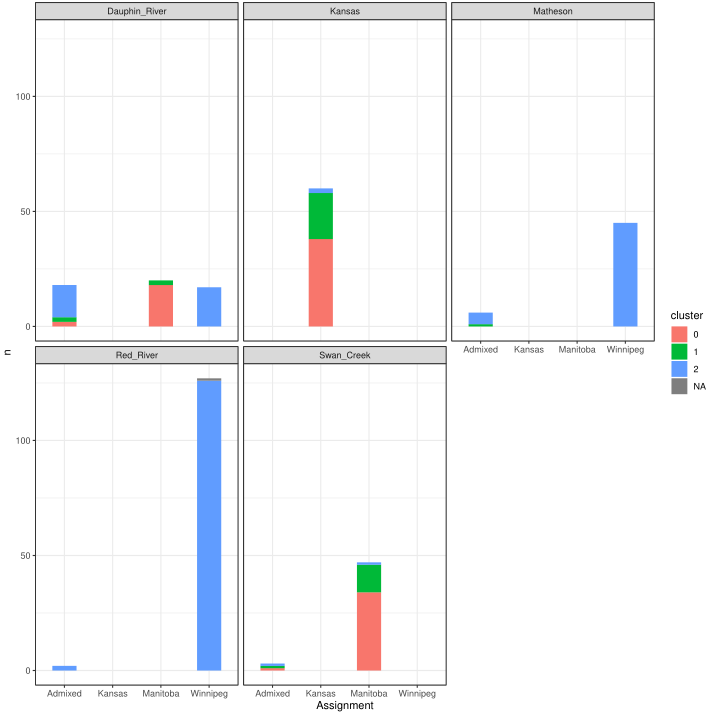

**Fig S9** Inversion frequency by assignment and site collected.

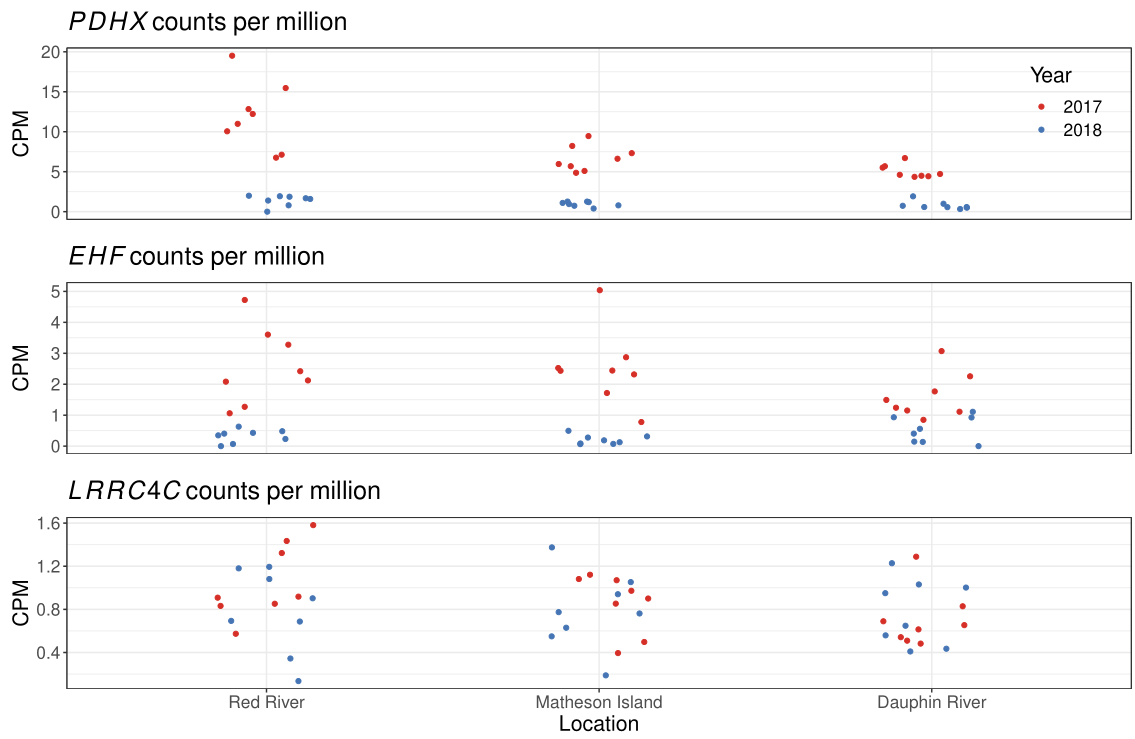

**Fig S10** Counts per million (CPM) for three genes within the putative inversion by site collected for walleye in 2017 and 2018.
